# Early developmental asymmetries in cell lineage trees in living individuals

**DOI:** 10.1101/2020.08.24.265751

**Authors:** Liana Fasching, Yeongjun Jang, Simone Tomasi, Jeremy Schreiner, Livia Tomasini, Melanie Brady, Taejeong Bae, Vivekananda Sarangi, Nikolaos Vasmatzis, Yifan Wang, Anna Szekely, Thomas V. Fernandez, James F. Leckman, Alexej Abyzov, Flora M. Vaccarino

**Author notes:** These authors contributed equally (Liana Fasching, Yeongjun Jang).

## Abstract

Post-zygotic mosaic mutations can be used to track cell lineages in humans. By using cell cloning and induced pluripotent cell lines, we analyzed early cell lineages in two living individuals (a patient and a control), and a postmortem human specimen. Of ten reconstructed post-zygotic divisions, none resulted in balanced contributions of daughter lineages to tissues. In both living individuals one of two lineages from the first cleavage was dominant across tissues, with 90% frequency in blood. We propose that the efficiency of DNA repair contributes to lineage imbalance. Allocation of lineages in postmortem brain correlated with anterior-posterior axis, associating lineage history with cell fate choices in embryos. Recurrence of germline variants as mosaic suggested that certain loci may be particularly susceptible to mutagenesis. We establish a minimally invasive framework for defining cell lineages in any living individual, which paves the way for studying their relevance in health and disease.

## Introduction

Organism development is hierarchical and involves several levels of cell organization beginning with the formation of the trophoblast (precursor of placenta) and inner cell mass (the embryo proper), continuing with formation of the embryonic disk, definition of left and right through formation of the primitive streak and then the segregation of embryonic cells into germ layers, i.e., the ectoderm, endoderm and mesoderm during gastrulation. Somatic mutations, which are those mutations that are generated after the formation of the zygote, can be used as permanent cellular markers to trace cell lineages and their spread throughout the human body (*1, 2*). It is estimated that 1 to 2 somatic SNVs are created per cell division during embryogenesis (*2, 3*), while in organogenesis the rate could be even higher (*2*).

Somatic mutations in cell lineages may contribute to differences in their characteristics. Cells in different lineages can have different rates of proliferation, senescence or death, all of which may affect the contribution of their cells to the adult body. In fact, it was observed that lineages contribute unequally to blood composition beginning from the first division of the zygote (*1*). Similarly, it is known that normal development may result in unequal characteristics of symmetrical organs, such as different volumes of left and right frontal and occipital regions of human brain (*4*). Furthermore, several pathological conditions exhibit asymmetrical manifestations such as motor symptoms in Parkinson Disease (*5*), hemimegaloencephaly and cortical dysplasia (*6*). One of the hypotheses that explain body asymmetries is through a causative role of somatic variants and their non-uniform contribution across the body (*7-9*), as proposed for Blaschko lines (*10*). However, the contribution of cell lineages to different portions of the human body is just beginning to be explored, and the timing and rules of their separation, spread and local expansion in organs are largely unknown.

To study this question, we reconstructed and analyzed early developmental cell lineages in a phenotypically normal living 66-year old female (NC0), a living 29-year old male patient with Tourette syndrome (LB), and in a post-mortem female fetus where we previously studied somatic mutations during neurogenesis (#316) (*2*).

## Results

For the two living individuals, NC0 and LB, we collected skin biopsies from two locations on left and right arms and one location on left and right thighs (**Fig. 1A**) and cultured fibroblasts from each biopsy. Multiple induced pluripotent stem cell (iPSC) lines were derived (a total of 74 for LB and 15 for NC0) and a selection of iPSC lines was sequenced from each fibroblast sample (**Table S1**). For the post-mortem fetal specimen 316, we re-analyzed eleven clones previously derived from telencephalic neuronal progenitors (*2*), and also sequenced 2 iPSC lines derived from dural fibroblasts. Most iPSC lines are clonal, and indeed, out of all iPSC lines, only one line showed evidence of being founded by two cells (**Fig. S1**). Cross comparing the genomes of iPSC lines/clones from the same person allowed us to discover the somatic mutations present in the founder cells of each clone (i.e. skin fibroblasts for LB and NC0, and brain progenitor cells for 316) (**Methods**). Sites with somatic variants were also analyzed in high coverage sequencing data for bulk blood, saliva, and urine from living individuals, and multiple brain regions and spleen from the fetal specimen, to determine their presence and precise allele frequency in those tissues.

**Figure 1.**
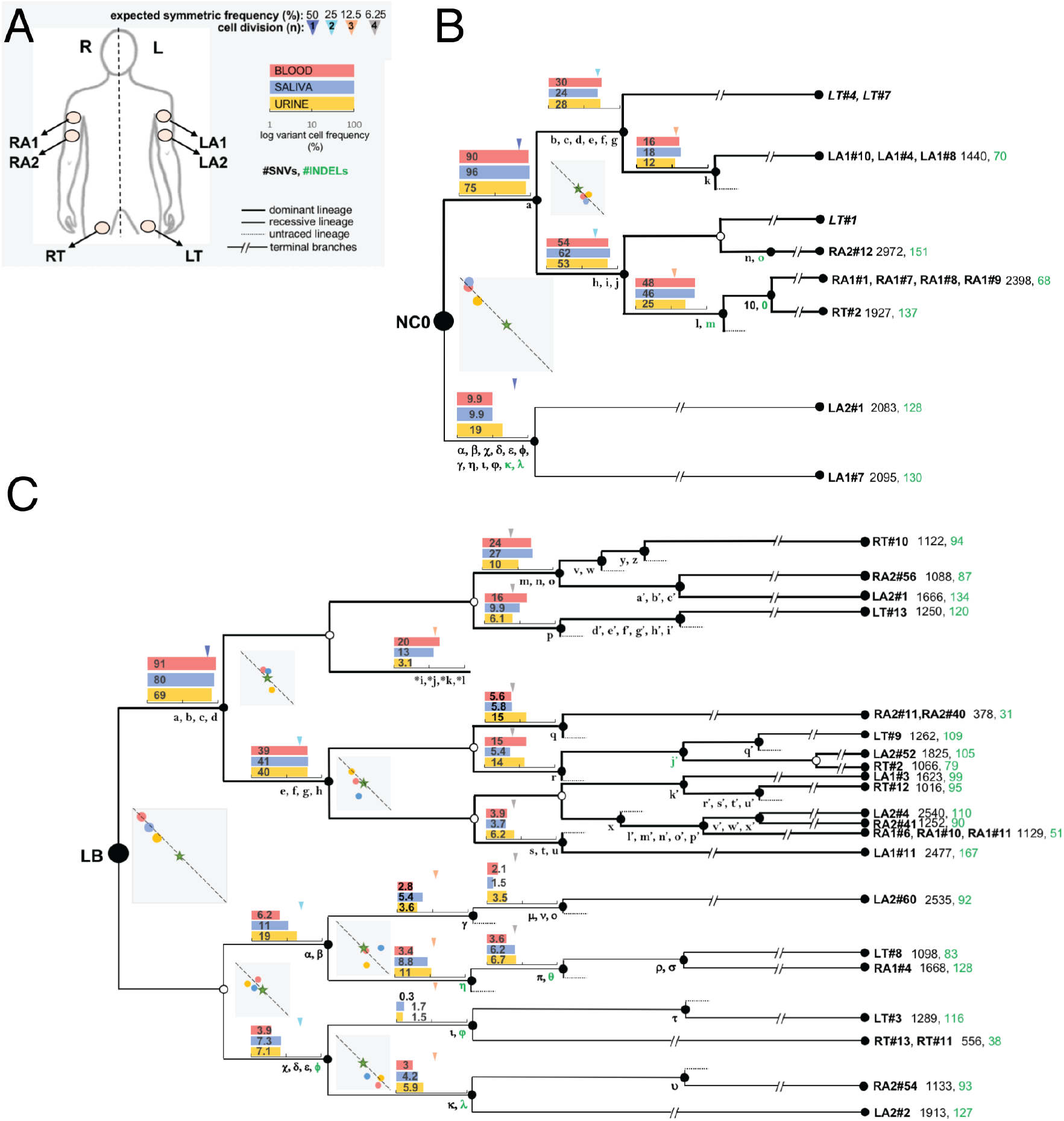
Reconstruction of early cell lineages in two individuals reveals lineage imbalance across tissues. **(A)** Outline showing location of the biopsies used to derive iPSC lines from skin fibroblasts. The iPSC lines (11 for NC0 and 25 for LB) used for making the trees are listed in (B) and (C) at the terminus of each branch, with 3 additional lines sequenced at shallow coverage in italics. **(B**,**C)** Early lineage trees. Cells are shown by circles with connecting edges representing parental relationships. Open circles mark ambiguous branching, i.e., where no new mosaic variant in the corresponding cell was found. Continuous lines depict pre-gastrulation lineages, with corresponding mosaic SNVs (black) and indels (green) denoted by Latin and Greek letters (see **Methods**). Broken lines depict terminal branches, with counts of somatic SNVs and indels present in each terminal branch. SNVs found in bulk tissues but not sampled by any of the iPSC lines are marked with asterisks. The average % variant cell frequency in blood, saliva and urine is shown in log scale by bar graphs next to the corresponding variants, with arrows indicating the expected cell frequency for a balanced lineage contribution. Squared plots show correlations of variant cell frequencies between the sister branches in blood (red), saliva (blue) and urine (yellow) (Y axis= upper branch; X axis= lower branch frequency), with stars indicating expected frequencies for balanced contributions. Frequencies of lineages in the two plots in NC0 and five plots in LB are consistent with the diagonal, suggesting that all progenies of these divisions are fully captured. Division at branch marked by e, f, g, h may not be fully captured, as frequencies in saliva are inconsistent with diagonal.

### Reconstructing early lineage trees

To reconstruct the early developmental cell lineages (starting from the first zygotic cleavage) of each individual, we selected somatic variants shared by clones/lines or by multiple bulk tissues (**Figs. 1, S2; Table S2; Methods**). To make branches in the lineage, we relied on variant sharing by lines/clones and on the rule that variants from consecutive cell divisions must be present in tissues at progressively decreasing frequencies. Therefore, variants within the same lineage, but having different frequencies in tissues are likely generated at different divisions. As expected from our selection criteria, all variants in the trees were shared by at least two tissues from different germ layers, i.e., mesoderm (fibroblasts, blood and spleen), ectoderm (brain and, in part, saliva), and endoderm (urine), and therefore arise in common progenitors of the three germ layers before gastrulation.

Occasionally, due to absence of variants, assigning lineages to a particular division was ambiguous, and multiple solutions in tree branching were possible (polytomy). In such cases, we chose the solution where sister branches had the most balanced frequencies in tissues, but alternative solutions were also recorded (**Figs. S2, S3**). To ensure completeness of reconstructed cell divisions, we compared frequencies of the reconstructed sister branches at each division in tissues. Normalized to the frequency of the mother cell, such frequencies should sum up to 100% in all tissues or, in other words, should be located on a *f*_*1*_*+f*_*2*_*=100%* line in a squared plot (**Fig. S4**). Within measurement errors, the frequencies of ten reconstructed divisions (2 in NC0, 5 in LB, and 3 in 316) fit the line, suggesting that these divisions are fully reconstructed (**Fig. 1, S2**).

### Imbalanced contribution of early cell lineages

For any given cell division, an equal 50%:50% representation of corresponding daughter lineages in the progeny would correspond to stars in the squared plots (**Fig. S4**). For most reconstructed divisions (starting from the first one) we observed imbalanced contributions of sister lineages in tissues, where dots in the squared plots are on the diagonal but away from the stars (**Fig. 1, S2**). In the two living individuals, the largest imbalance was observed for the first two blastomeres revealing the presence of a dominant and a recessive cell lineage (**Figs. 1, 2A**). In both individuals, one of the blastomeres generated 70%-90% of cells in tissues, while the second blastomere generated 30%-10% of the cells in the final tissues. The largest imbalance was present in blood, with a contribution ratio of 90:10 from dominant:recessive blastomeres in both NC0 and LB, while the smallest difference was in urine, with roughly 70:30/80:20 contribution ratio in the 2 individuals; urine showed the least deviation also in subsequent cell divisions (**Fig. S5**). The ratio in saliva was roughly the same as in blood for NC0 and was somewhere in-between those in blood and urine for LB (**Fig. 2A**). Saliva in adults is estimated to consist of about 50%-70% of leukocytes (mesoderm) and 50%-30% of epithelial cells (ectoderm) (*11, 12*). Hence for LB the contribution ratio for ectoderm should be somewhat lower and probably closer to that present in urine. In the postmortem individual no marked imbalance of the first two blastomeres existed, though one of the two alternative trees is consistent with a 90:10 imbalance in the first division (**Fig. S2**). The ambiguity also existed for LB, but the alternative trees only increase the imbalance (**Fig. S3**).

**Figure 2.**
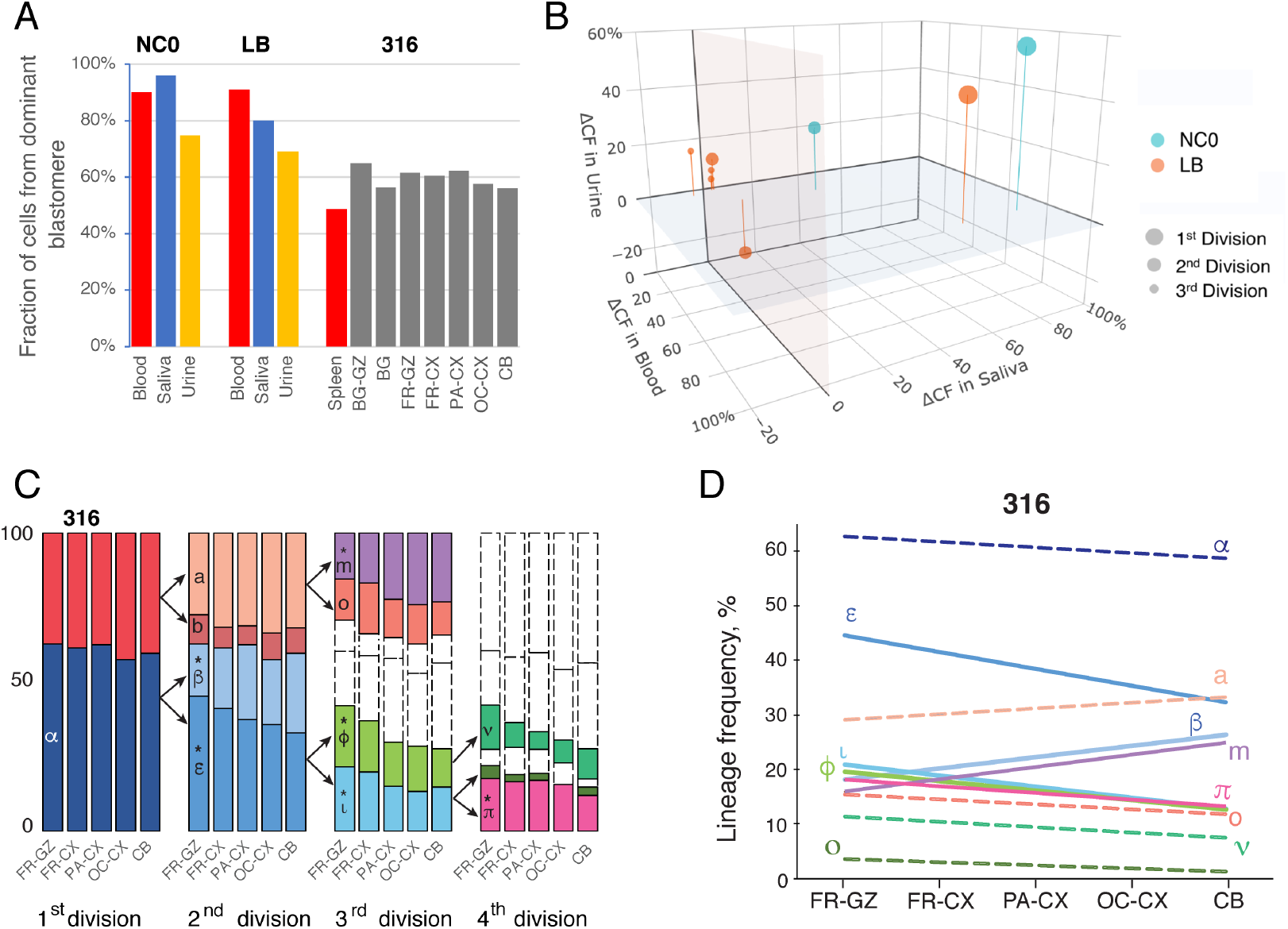
(**A)**, fraction of cells contributed from the dominant blastomeres in the first cell division to various tissues in NC0, LB and 316. (**B)**, difference of sister lineages contribution to blood, saliva and urine for fully reconstructed cell divisions in NC0 and LB individuals. (**C)**, percent of lineage contribution to different brain regions at each cell division in fetal specimen 316. FR-GZ, frontal germinal zone; FR-CX, PA-CX and OC-CX are frontal, parietal and occipital cortex, respectively. Frequency of lineages is represented by bars of different colors, with letters denoting representative variants for each lineage (**Fig. S2**). The arrows between divisions indicate the correspondence of a mother cell to respective daughter lineages. (**D)**, linear regression of lineage frequency across brain regions. Significant correlations (p-value < 0.05 by Spearman rank-order) are shown by solid lines and are marked by stars in C, while non-significant correlations are shown by dashed lines. Lineages with marginal significance (p-value < 0.1) are marked by variants ‘α’, ‘a’ and ‘o’. Regressions are not shown for recessive lineage from first division and lineage marked by ‘b’ variant from second division, as their frequencies are dependent from frequencies of the other shown lineages for these divisions

Higher prevalence of one sister lineage over the other one was typically consistent across tissues for all divisions (**Fig. 2B** and **S5)**, suggesting consistency of early lineage allocations across germ layers and the body. Imbalance in early lineage allocation may result from intrinsic properties of the founder cells, and/or be the result of selection, which could be different in different tissues. Here, we noticed that in LB (**Fig. 1**) there is a higher fraction of indels among variants of the recessive lineage as opposed to the dominant lineage (5 vs 1, p-value = 0.03 by Fisher’s exact test) (see **Methods**). Because of a tree with fewer branches, such indel imbalance could not be verified for NC0. But interestingly, the recessive lineage in this individual had 10 SNVs and two indels in the cell at its origin (compared to only one SNV in the dominant lineage) (**Fig. 1**). Consistent with that, one of the alternative lineage trees for 316 with the largest imbalance in sister lineage allocation to tissues also had a disproportionally large count of variants (i.e., 11 vs 0) in the recessive as compared to the dominant lineage (**Fig. S2C**). Based on these observations, we hypothesize that the efficiency of DNA repair in the early divisions of the human embryo is a factor contributing to lineage imbalance.

### Lineage distribution in relation to anterior-posterior axis

The available data allowed us to gain insight into lineages from the 1^st^ to 4^th^ cell division (first week of development), which corresponds to the human embryo pre-implantation stage. It is debated whether at this stage there are clear laterality or anterior-posterior (A-P) bias in lineage allocation, although apical-basal polarity, which might anticipate other type of asymmetries, is present at the 8-cell stage in mouse embryos (*13*). Gastrulation, occurring approximately at the beginning of the third week of human embryonic development, is when antero-posterior and left-right asymmetry is thought to occur in development, through the formation of the primitive streak. If this is true, there should be no systematic bias in pre-gastrulation lineages allocation with respect to body axes. Using genotyping of mosaic SNVs from high depth resequencing, we inferred frequencies of early lineages across multiple brain regions of specimen 316. Lineage frequencies in tissues are not independent, e.g., frequency of the recessive lineage is strictly bound to frequency of the dominant lineage. We therefore analyzed 11 lineages with independent frequencies (see **Methods**). Six lineages showed significant correlations (p-value < 0.05 by Spearman rank-order) and three more marginally significant correlations (p-value < 0.1) between their frequencies in dorsal brain regions and the arrangement of these regions along the A-P axis, as exemplified by the order FR-GZ => FR-CX => PA-CX => OC-CX => CB (**Fig. 2C**,**D**). The two remaining lineages had the lowest frequencies, consistent with significance not being detected due to larger errors in measuring their tissue frequencies. This analysis reveals association of early lineage allocation with respect to the A-P axis in brain.

### Recurrence of population and mosaic variants

In the LB patient, two early mosaic SNVs used for reconstruction of the lineage tree (SNP β and ε, see **Table 2**) matched to germline single-nucleotide polymorphisms (SNPs) in his mother (**Figs. 3, S6**). Both SNPs were also present in the catalogues of germline variants by gnomAD (one with frequencies in human population of more than 0.001). Examination of the SNV sites revealed nearly 50% variant allele frequency (VAF) in all iPSC lines where the SNVs were called, 0% (or near 0% in one line) VAF in all iPSC lines where the SNVs were not called, and intermediate VAF in blood, saliva, and urine (**Table S2**). We found no deletion or loss-of-heterozygosity in the lines missing the variants, which could have accounted for the variants being inherited but absent in some lines. Furthermore, population-based phasing of germline variants in the patient and his parents was consistent with the scenario that haplotypes with the matching SNPs in LB’s mother were not inherited by LB. Ultimately, physical read backed phasing confirmed the nonhereditary nature of the mosaic SNV β in LB (**Fig. 3**). We, therefore, concluded that these SNVs are bona fide mosaic variants recurrent of known population variants. Sequence context aware simulation (**Methods**) of a random match of mosaic and germline variants suggested that a count of two recurrent SNVs is unlikely to happen by chance (p-value = 0.006). In addition, we also found that 3, 1, and 3 of early mosaic SNVs in LB, NC0 and 316, respectively, match known frequent (>0.001 allele frequency) population variants in gnomAD (**Table 2**), which is unlikely to happen by chance according to the same simulation (p-value = 0.03). In sum, we detect an unexpectedly high recurrence of population variants as post-zygotic variants during embryonic development. Interestingly, SNV β discussed above occurs at the boundary of two homopolymers (T)_12_(A)_6_. Homopolymers are known to be prone to expansion and contraction (*14*). It is therefore possible that the SNVs is the results of T-homopolymer contraction and A-homopolymer expansion, i.e., due to overlap of two indels: TT>T and A>AA. This possibility is also consistent with the above hypothesis that recessive lineages have a higher fraction of indels, as the discussed SNVs is found in branches within the recessive lineage (**Fig. 1**).

**Figure 3.**
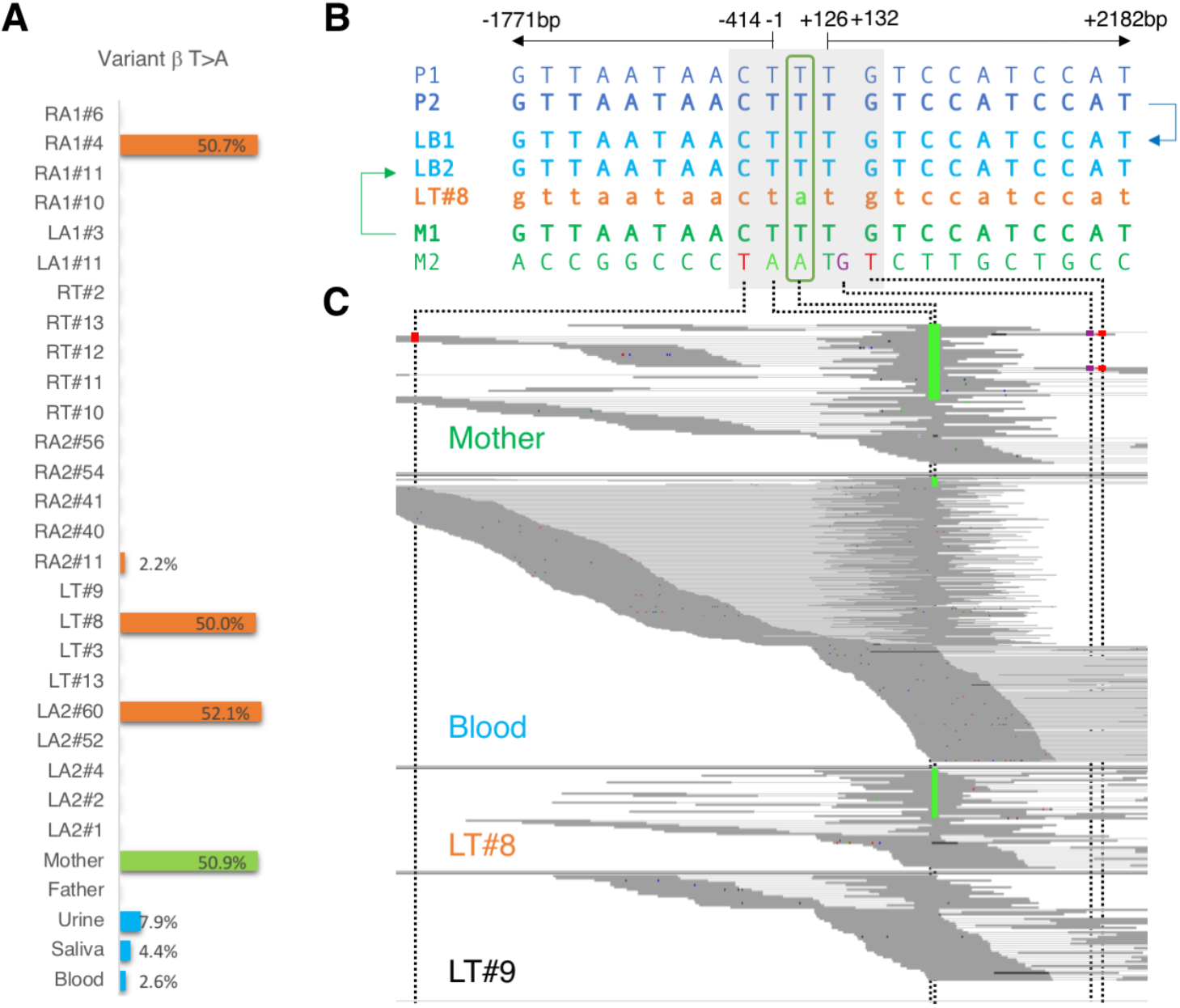
Recurrence of a germline SNP in LB’s mother as a mosaic SNV in LB. (**A**) VAF of the T>A somatic SNV variant across iPSC lines (orange), blood in parents (green), and urine, saliva and blood in the patient (blue). (**B**) Schematics of germline haplotype inheritance based on population-based SNP phasing for LB (LB1, LB2), father (P1, P2), and mother (M1, M2). Ten variable positions in parents downstream and upstream from the SNV are shown. (**C**) Read level evidence for the maternal haplotype with the variant not being inherited by the patient. Every row has a single connected (light grey lines) pair of reads (dark grey lines). The SNP in the mother is in phase with 4 nearby variants, none of which is present in the patient.

## Discussion

Here we developed and successfully applied a minimally invasive framework for studying early lineages and their imbalance in living human individuals, which consists of two components: 1) derivation and analysis of genomes of clonal iPSC lines to discover somatic mutations and conduct lineage reconstruction; and 2) analysis of variant/lineage distribution in bulk tissues such as saliva, blood and urine to establish their hierarchy in development. In future studies iPSC lines from skin fibroblasts can be complemented by iPSC lines from other easily accessible cells: epithelial cells from urine (*15*), erythroblast from blood (*16*), and keratinocytes from skin (*17*), expanding the diversity of sampling of early lineages. Analysis of DNA from bulk blood, saliva, and urine can be complemented by the analysis of DNA from buccal swabs, multiple skin regions, feces, vaginal cells and sperm. Therefore, our study paves the way for comprehensive and large-scale analyses of early lineages and understanding their role in human health and disease.

We show the existence of dominant and recessive lineages starting from the first cell division of the human embryo, demonstrating an imbalanced lineage allocation across tissues of the human body on a global scale. Our study supports previous suggestions of an imbalanced lineage contribution to tissues in the human body after the first cleavage (*1, 3*); however, our results point to a generally higher imbalance of 90:10 vs 2:1 suggested previously. Only minor deviations from the general trend in lineage allocation at each division were observed within each germ layer, implying that the imbalance was established by a general mechanism in the original lineage precursor cells rather than selective processes within each tissue compartment. In brain, sister lineage allocation among the first 4 divisions showed a correlation with A-P axis development. The ordered pattern of lineage distribution along the A-P axis across several brain regions argues against this phenomenon being regulated by selective pressure exerted on a particular lineage in a particular tissue compartment, and suggests that intrinsic phenomena in lineage founder cells may bias their daughter’s allocation according to an A-P axial rule at very early stages of human embryonic development. Cell intrinsic differences underlying differential lineage contribution to tissues are likely rooted in gene regulation or epigenetic phenomena in the lineage precursor cells (e.g., unequal repartition of cellular determinants in the progeny) potentially affecting different rates of proliferation and/or survival of daughter cells (*13*).

We observed an excess of indels in the recessive versus the dominant lineage, and we hypothesize that one factor contributing to this imbalance is the efficiency of DNA repair. Indels can be created from polymerase slippage and from faulty mismatch repair (*18*), while in development most SNVs arise from spontaneous deamination of 5-methylcytosine (*19*). Additional time spent by a cell on DNA repair may decrease proliferation rate leading to lower contribution to tissues (**Fig. 4A**). This line of reasoning is applicable to patient LB. For normal individual NC0 and the fetal specimen, no excess of indels, but rather an increase in SNV burden in the founder cell of the recessive lineage was observed. We hypothesize that this increased burden could be also caused by deficient DNA repair generating genomic instability in the recessive lineage, a phenomenon hypothesized to exist *in vivo* based on analysis of *in vitro* fertilized embryos (*20*). The instability may result in multiple consecutive cleavages giving only one viable daughter cell, and thus leading to accumulation of point mutations from such cleavages in the viable cell; these mutations would retrospectively seem to occur from a single division (**Fig. 4B**). An alternative hypothesis to the importance of DNA repair efficiency could be that cell fate determination happens already at the first cleavage, where one of the created blastomeres commits mostly to the extra-embryonic lineages and its progenies migrate away from the embryo, while the other blastomere mostly commits to inner cell mass and becomes the dominant lineage. Although debated (*13*), such a possibility was previously proposed for mammalian embryos (*21*). In such case, cells segregation into trophoblast would play the same role as non-viable cells in the scenario discussed above, i.e., they will not be present in the adult body (**Fig. 4B**). Thus, lineage studies can shed light onto mechanisms that regulate cell fate decisions during development as well as representation of different lineages in tissues of any living individual and across the human population.

**Figure 4.**
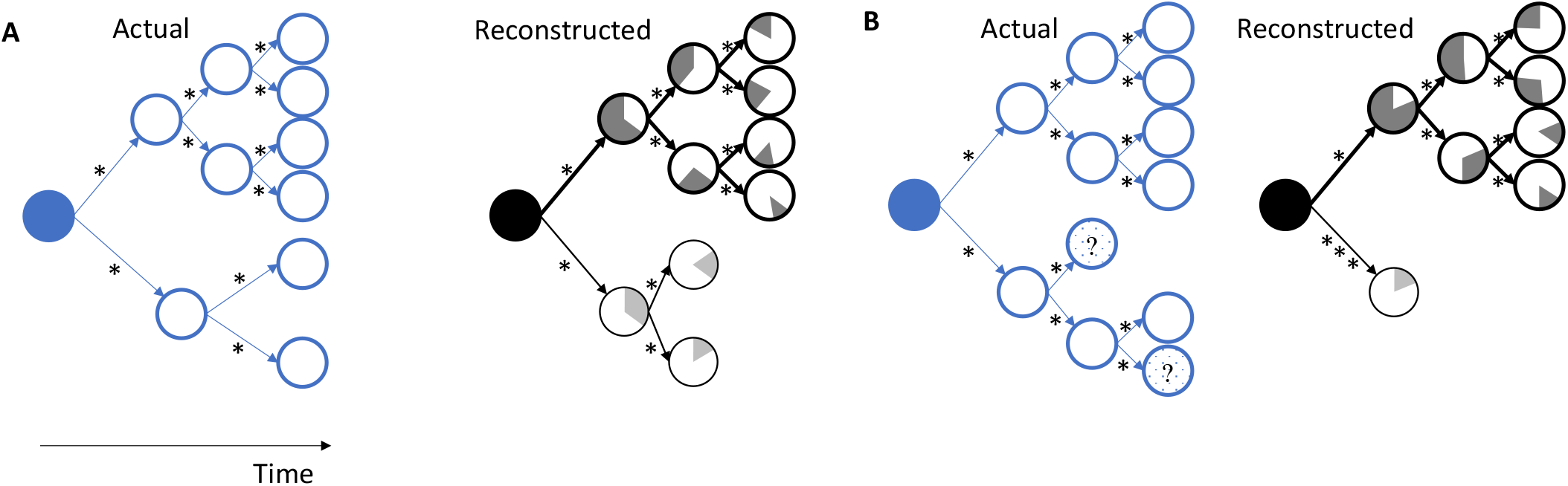
Hypothetical scheme of lineage imbalance in tissues of the adult body arising from early developmental events. (**A**) Cells of the bottom lineage divide more slowly compared to the top lineage, making the bottom lineage recessive (i.e., underrepresented) in adult tissues. Mutation burden in branches of the recessive lineage could be the same as for the dominant lineage. (**B**) Cells in the bottom lineage have more progenies that are not viable or chose more frequently an extra-embryonic fate (dashed circles with question marks) compared to the top lineage, making the bottom lineage recessive (i.e., underrepresented) in adult tissues. Reconstructed branches in the recessive lineage will have higher mutation burden than in dominant lineage. ‘*’ represents a mutation. Shading within cells represent mutation frequency in adult tissue (i.e., blood).

## Acknowledgments

We are most grateful for LB and his parents as well as NC0 for their willingness to participate in this study. We acknowledge the Yale Center for Clinical Investigation for clinical support in obtaining the biopsy specimens. We thank Dr. Caihong Qiu and Jason Thomson of the Yale Stem Cell Center Core Services for the generation of iPSC lines. We thank BGI Americas Corporation for library preparation and deep sequencing. We thank the Keck DNA Sequencing Facility at Yale for their assistance with DNA sequencing service. We thank Drs. Jessica Mariani and Soraya Scuderi for the generation and amplification of brain neurosphere clones. We wish to acknowledge members of the Brain Somatic Mosaicism Network (BSMN) for helpful comments and discussions. This work was funded by the National Institute of Mental Health (grants R01 MH100914, U01 MH106876).

## Methods

### Sample collection

Informed consent was obtained from each subject according to the regulations of the Institutional Review Board and Yale Center for Clinical Investigation at Yale University. Skin biopsies (approx. 3mm^3^) were collected respectively from the inside of the upper left and right arms (two locations about 5 cm apart and corresponding to different dermatomes) and upper left and right thighs and cultured separately in fibroblast culture media (DMEM high glucose, FBS, L-glutamine, N.E. amino acids, Pen/Strep; all Invitrogen). Fibroblasts attached, proliferated and were passaged twice before DNA was extracted (DNeasy Blood and Tissue Kit; Qiagen according to manufacturer recommendations).

Saliva DNA was collected and purified using the Oragene-Discover kit (DNA Genotek) following the manufacturer instructions. Saliva DNA was extracted using DNeasy Blood and Tissue kit (Qiagen) with the following modifications: 5 ml AL-buffer and 200 μl Proteinase K were added to collected saliva in buffer and incubated at 56°C for 30 minutes. RNA was digested using 20ul RNAse A (Qiagen) for 5 minutes and DNA was extracted using 4 extraction columns in parallel to optimize the yield.

For the purification of DNA from urine, urine was collected into 50ml falcon tubes and kept on ice during transport. Cells were pelleted by centrifugation (400g for 10 minutes at RT). Supernatant was aspirated and cells were resuspended in 500ul PBS and 1.5ml of lysis buffer (DNeasy Blood and Tissue kit (Qiagen) per each 50ml falcon tube. 100ul of Proteinase K were added to each tube, vortexed and incubated at 56°C for 25 min. 10ul of RNAse A were added to each sample and incubated for 5 minutes. DNA was extracted using the DNeasy Blood and Tissue kit (Qiagen).

### Blood collection and DNA purification

For LB, his mother and father and NC0 10-15 mL of blood was collected using BD Vacutainer ACD tubes. DNA was extracted using the Gentra Puregene Blood Kit (Qiagen) following standard manufacturer protocols.

### iPSC line derivation

iPSC lines were derived using the Epi5 Episomal iPSC Reprogramming Kit (Invitrogen catalog A15960) which delivers the five reprogramming factors Oct4, Sox2, Klf4, L-Myc, and Lin28. iPSC lines were propagated using mTeSR1 media (Stem Cell Technologies). Genomic DNA was extracted at passage six, using QIamp DNA Minikit (Qiagen) following the manufacturer instructions.

### Sequencing of bulk tissues and iPSC lines

Whole genome sequencing (WGS) of bulk DNA samples from living individuals was conducted at 200X, while WGS of iPSC lines was typically conducted at 30X, with some lines sequenced at 5X. All sequencing was conducted at BGI using with 2×100 bp paired reads. For all but urine samples sequencing library preparation was PCR-free.

### Calling mosaic variants in iPSC lines and clones

Mosaic SNVs and indels were called from an exhaustive all-2-all comparison of all lines/clones for each individual. For each pairwise comparison of lines/clones we used calls made by both MuTect2 (*22*) and Strelka2 (*23*). Previously we described an approach for selecting bona fide mosaic variants from such comparison (*2*). Conceptually, the approach selects calls that are consistently made when comparing lines from a set A to lines from a complementary set B (sets A and B account for all lines/clones in an individual). Here we extended the approach to select calls when they are found in most lines/clones (i.e., when set A is much larger than set B) and thereby resemble germline variants. Calls were required to have 35% allele frequency in line/clones. Additionally, only indels shorter than 10 bp were retained to reduce false positive rate. The filtering tool is freely available https://github.com/abyzovlab/all2. To protect individual’s privacy, the coordinates of discovered variants were de-identified in the manuscript. Full information is available from the database (see **Data access**).

### Calling mosaic variants in bulk tissue DNA

To call variants from bulk tissue we followed best practice development by the Brain Somatic Mosaicism Network (unpublished). Briefly, calling consisted of the following steps: 1) call variants with GATK using a ploidy setting of 50; 2) eliminate calls in inaccessible genomic regions according to the 1000 Genomes mappability mask; 3) discard germline variants that have a population allele frequency of >0.001 in gnomAD catalogue; 4) eliminate calls consistent with ∼50% VAFs by a binomial test significance of 10^−6^; 5) eliminate calls in genomic regions exhibiting copy number gains; 6) mandate that candidate mosaic SNVs have at least 5 independent supporting reads that have minimum values of 20 for mapping and 20 base quality; 7) identify and eliminate false-positive calls using samples in the 1000 Genomes Project as a panel of normal filter; 8) filter calls using MosaicForecast tool (*24*).

### Lineage tree reconstruction in LB

We first sequenced six iPSC lines to 30X coverage and discovered mosaic variants in those six lines using the all-2-all exhaustive comparison described above. We next constructed the early lineage tree and selected 14 variants marking branches of the tree. To avoid sequencing redundant iPSC lines derived from the same fibroblast clone in the skin, we then genotyped all the iPSC lines for the presence of those fourteen mosaic variants using amplicon-seq (see below). Based on genotyping, the lines were assigned to existing and new branches in the tree. This allowed us to select from each biopsy, one line likely contributing a new branch to the lineage tree for further analysis, i.e., nineteen additional lines, that were sequenced at 30X coverage. We then called mosaic variants using the all-2-all exhaustive comparison from all 25 of the sequenced iPSC lines (see above; **Table S2**). For each called mosaic variant we then estimated its VAF in bulk tissues (saliva, blood, urine) by using the mpileup function of samtools requiring minimum values of 20 for mapping and 20 base quality. Estimates of VAF for low frequency variants are less reliable and to select most confident sites we paralleled VAF estimated by a different approach. Namely, we re-called variants in the bulk using GATK with ploidy 100 and extracted from the output the number of reads supporting reference and alternative alleles. Such ploidy value allows discovering SNVs with VAF below 1% and, combined with 200X coverage of bulk allows for VAF estimation for variants with at least 2 supporting reads (i.e., for over 90% of variants with actual VAF of 2%). Out of the called mosaic variants we chose a subset of early (pre-gastrulation) variants for the lineage tree construction using the following criteria: (1) a mosaic variant (SNV or indel) is shared by at least two iPSC lines from different biopsies; or (2) a mosaic SNV has at least 2% VAF in at least one bulk tissue and at least one supporting read in another bulk tissue.

In addition to mosaic variants discovered in iPSC lines, discovery in bulk DNA from blood, saliva and urine revealed 4 high VAF mosaic SNVs not sampled in any of the iPSC lines. We placed them in a separate branch to complement the branches reconstructed from analysis of iPSC lines. The criterion we used is that the sum of VAFs in this newly created branch with its sister branch should be roughly equal to the parental branch (**Fig. S4**). One SNV discovered in LA2#4 conflicted with the tree (had VAF in bulk tissues higher than variants in preceding branches) and was not used in the tree construction (**Table S2**).

### Lineage tree reconstruction in NC0

For each of two biopsies in the left and right arm we sequenced four iPSC lines to 30X coverage. For each one of the other four biopsies, we sequenced one iPSC line to 30X coverage. The line from left thigh was excluded from the analysis because of likely originating from two cells (**Fig. S1**). Therefore, we additionally sequenced three iPSC lines from left thigh to 5X coverage. We then called mosaic variants from exhaustive comparison of the 11 iPSC lines with 30X coverage. The lineage tree was constructed as described for LB. One SNV was excluded from tree construction as likely germline variant. The exclusion was based on the following observations: 1) the variants had VAF above 45% in blood, saliva, and urine; 2) the variant is present in human population with allele frequency of over 10%; 3) the variant was present in all but three redundant (i.e., sharing almost all of their variants) iPSC lines (LA1#4, LA1#8, LA1#10), consistent with it being inhered but lost in fibroblasts from the LA1 biopsy; 4) the variant conflicts with the branches in the tree (**Table S2**). Three additional lines with 5X coverage were placed into the tree by genotyping variants in these lines from the constructed tree. Mosaic variant ‘a’ with the highest VAF was validated by Sanger sequencing (**Fig. S7**).

### Lineage tree reconstruction in 316

We previously called mosaic variants in human brain progenitor clones of the post-mortem fetal specimen 316 and constructed an initial tree of early lineages assuming equal contribution of sister lineages (*2*). Given the evidence of imbalanced lineage contribution in LB and NC0 we removed this assumption. Additionally, using the extended filtering approach described above, we re-called and filtered mosaic variants in previously obtained WGS data for clones and added new data for two iPSC lines derived from skull fibroblasts of the same fetal specimen. Compared to the previous call set, we found 3 additional SNVs shared by multiple clones/lines, all of which were at high frequency in brain regions and spleen (VAF of 10%-35%), allowing to resolve the common ancestry of two pairs of lineages (i.e., making two unambiguous branches) (**Fig. S2**). Mosaic variant ‘α’ with the highest VAF was validated by Sanger sequencing (**Fig. S8**). We also found 3 indels (indels were not analyzed previously) shared by multiple lines/clone, which all supported branches based on SNVs. Previously, we used a capture-seq validation approach to determine the frequency for discovered mosaic variants across brain regions. Using that data, we previously omitted from lineage reconstruction variants for which we could not determine precise VAF in tissue. Because of using 125 bp baits, capture-seq has limited power of enriching for sequence in repeats of comparable length. Contrary to that, WGS is powered to call variants in repeats of length up to the fragment size of DNA being sequenced, i.e., up to 300-500 bps. Following this consideration, we now included all SNVs shared by at least 2 clones/lines variants (7 additional SNVs) for tree reconstruction. These additional variants allowed resolving a conflict that existed in the previous tree by adding a lineage split (i.e., an additional branch) (**Fig. S2**). Following the same criteria as for LB and NC0 we also used mosaic SNV with at least 2% VAF in at least one bulk tissue and at least one supporting read in some other bulk tissue for lineage reconstruction. The lineage tree was constructed as described for LB and NC0.

### Comparing fractions of indels in lineages in LB

Indels are known to be more challenging for discovery and genotyping than SNVs. Because of that, we only used indels that were discovered in at least 2 iPSC lines from different biopsies to reconstruct the lineage tree. For consistency, we applied the same considerations to SNVs when comparing fractions of indels in dominant and recessive lineages. We did not count variants from the first division since as at this point the dominant/recessive lineages did not exist. The dominant lineage had 22 variants shared by iPSC from different biopsies, which included only 1 indel. The recessive lineage had 15 shared variants with 5 indels. The proportion of indels was significant by Fisher’s exact text (p-value = 0.03).

### Analysis of lineage distribution in fetal brain

There are total 13 reconstructed lineages in postmortem specimen 316 (**Fig. 2C**). Of them, frequencies of two lineages are dependent on the frequencies of other lineages: frequency of recessive lineage is dependent on the frequency of dominant lineage and frequency of lineage ‘b’ is dependent on the frequencies of lineages ‘a’, ‘α’, and ‘ε’. To reduce redundancy, those two lineages (recessive and ‘b’) were not included in the analysis.

### Sequence context aware simulation

To assess the chance of the discovered mosaic SNVs to match germline SNPs across the genome, we applied a sequence content aware simulation (i.e., we considered the tri-nucleotide sequence around the variants). In each round of simulation, we randomly scattered all the identified mosaic SNVs across positions in the genome with the same tri-nucleotide sequence. We only considered SNVs in P-bases and scattered them across P-bases of the 1000 Genomes Project accessibility mask. We then counted the fraction of SNVs matching to germline SNP in parents (for LB) or in gnomAD with population frequency of at least 0.1% (for all 3 individuals). We then repeated this process 1,000,000 times, building the null distribution of the number of mosaic SNVs matching to germline SNPs. We then calculated p-value of finding exact or larger numbers of observed recurrent mosaic SNVs using these distributions.

### Amplicon-seq and Sanger-seq validations and processing

Amplicon-seq was used to genotype 14 SNVs in iPSC lines, blood, saliva, and urine of LB. Primers (**Table S3**) were designed by selecting a DNA template of 800 nucleotides which contains the candidate SNV (SNV ± 400bp), with an amplicon size between 200-450bp. For PCR amplification, we used Phusion High-Fidelity DNA polymerase (Thermo Fisher Scientific) to minimize the polymerase error rate; optimal annealing temperature was defined by the Tm calculator tool available on the Thermo Fisher Scientific website and validated by PCR. Primer specificity was initially determined *in silico* with UCSC Genome Browser (http://genome.ucsc.edu/index.html), and then confirmed on the 2% agarose gel by the presence of a unique PCR product of the expected size. Amplicon DNA was purified using the QIAquick PCR Purification Kit (Qiagen). PCR products from the same sample were pooled and samples were submitted for sequencing under following conditions: MiSeq paired-end, 250bp.

SNVs “a” discovered in NC0 (**Fig. S7**) and “α” discovered in 316 (**Fig. S8**) were confirmed by Sanger sequencing. Primers for Sanger sequencing were designed as described for amplicon-seq and PCR products were visualized on 2% agarose gels. 40-50ng of amplified DNA per sample were submitted for sequencing.

## Data access

Identified variants and full supplementary tables have been deposited to the NDA database and associated with the study #930 (DOI:10.15154/1519174). Read level data are being uploaded to the same database and will be linked to the study upon completion of the deposition.

